# The genetic networks of regeneration, cell plasticity, and longevity of the Immortal Jellyfish *Turritopsis dohrnii* (Cnidaria, Hydrozoa)

**DOI:** 10.1101/2025.07.02.660568

**Authors:** Yui Matsumoto, Maria Pia Miglietta

## Abstract

When medusae of *Turritopsis dohrnii* are damaged, wounded, or exposed to otherwise lethal conditions, they revert to an earlier life cycle stage (the polyp) through an intermediate and transient benthic stage, the cyst, thus effectively escaping death. By employing a super transcriptome approach, we profile how the expression putative homologs of genes involved in regeneration, pluripotency, and longevity, change throughout life cycle stages of *T. dorhnii.* We follow the expression of putative homologs of Sirtuins, factors that control telomere maintenance, heat shock proteins (HSPs), and the Yamanaka transcription factor families (POU, Sox, Klf, Myc). We show that during its life cycle reversal, *T. dohrnii* manipulates genetic networks of high relevance in biomedical studies in mammals, such as SIRT3, POU factors, RTEL1, and HSP70/90. Our data showcase *T. dohrnii* as an *in vivo* research system that can contribute to understanding the genetic networks that regulate cell programming and ontogeny reversal.

## Introduction

Regeneration is the process by which organisms replace or restore lost or damaged tissues, organs, or body parts. It involves complex cellular processes, including stem cell activation and cell differentiation, and generally declines with age. Certain organisms exhibit remarkable regenerative abilities and potential pathways to biological immortality. Among these, the Cnidarian *Turritopsis dohrnii*, commonly known as the immortal jellyfish, stands out as a prime example of regeneration and longevity. Described from the Mediterranean Sea in 1883 (Weismann), *T. dohrnii*’s unique life cycle was recognized only in 1992 (Bavestrello et al. 1992, then redescribed in Martell et. al. 2016). When faced with stressful conditions such as lack of food, sudden changes in salinity, or physical damage, the medusae of *T. dohrnii* avoid death by reverting to the polyp stage (Bavestrello et al. 1992; Piraino et al. 1996, 2004; Miglietta and Lessios 2008). During the reversal, a series of events is recognizable: the gelatinous layer between ectoderm and mesoderm thins out, swimming is impaired, and the jellyfish settles on the bottom and becomes a cyst-like mass of cells with no medusa features, which will start showing polyp features in the following 24-72 hours. The process is a true metamorphosis, albeit in the opposite direction of the normal developmental cycle, and it has earned *T. dohrnii* the name of “immortal medusa”.

*Turritopsis dohrnii* challenges basic aspects of aging theories. For example, the onset of sexual maturation/reproduction is regarded as the point of no return in the ontogenetic sequence in living organisms, ultimately leading to death (Stearns, 1992). In *T. dohrnii,* ontogeny reversal can occur at any stage of medusa growth, including the adult stage. In *T. dohrnii,* senescence induces metamorphosis (Piraino et al. 1996; Schmich et al 2007). The life cycle reversal of *T. dohrnii* is difficult to frame in the typical regeneration landscape. It can be induced by physical wounding but can also be caused by other stressors, including starvation or exposure to heat. Moreover, the nature of the transition from adult medusa to polyp is hardly understood. Polyps and medusae differ in anatomy and cell types. Some cell types (i.e., mononucleated striated muscle cells, nerve cells, and sensory cells) are present in the medusa only. In contrast, other cell types (i.e., ectodermal cells producing chitin) are present in the polyps only. Medusae and polyps also contain interstitial cells (I-cells), cyclic embryonic cells that divide continuously and can differentiate into several cell types (Tardent 1978, Müller et al. 2004, Frank et al. 2009). Surprisingly, although I-cell proliferation occurs during the reversal, it does not play a crucial role, as portions of the jellyfish containing no I-cells can still revert to the polyp stage (Piraino et al. 1996). Given the limited role of I-cells, and the fact that medusae and polyps contain different cell types, the transformation from jellyfish to polyp has been hypothesized to involve cellular transdifferentiation (reviewed by Okada, 1991; Schmid, 1992) that occurs in a cyst-like stage, during which all the medusa characters disappear, and the first polyp characters appear in 24-36 hours. (Piraino et al. 1996). However, no cell tracking or single-cell transcriptomics has been performed to confirm the occurrence of cellular transdifferentiation. Regardless, *T. dohrnii* has a unique potential for rejuvenation. Because its life cycle reversal can be induced under controlled laboratory conditions, this species represents an unparalleled model for investigating aging, stress response, and regeneration.

Recent transcriptome analyses of *Turritopsis dohrnii* life cycle identified lifespan and aging, response to DNA repair and damage, and cell specification and differentiation, as gene ontology (GO) categories highly expressed in the cyst (Matsumoto et al. 2019; Matsumoto and Miglietta 2021). However, the role that genes known to directly influence regenerative capabilities, cell pluripotency, and the progression of aging in established model systems play in *T. dorhnii*’s life cycle has never been investigated. In this paper, using a supertranscriptome approach, we investigate how the expression of selected genes canonically associated with these biological processes of regeneration, such as Sirtuin proteins (Amano and Sahin 2019), telomerase (Flores and Blasco 2010), heat shock proteins (HSPs) (Tower 2011), and the Yamanaka transcription factor families (*Oct, Sox, Klf, and Myc*) (Takahashi et al. 2007), change throughout *T. dohrnii’s* reverse development.

## Methods

### Construction of Super-Transcripts and BLAST protein assignment

Super-transcripts (Davidson et al., 2017) were constructed from published RNA-sequencing (RNA-seq) libraries of life cycle stages of *T. dohrnii* (colonial polyp, medusa, cyst, reversed polyp) from Bocas del Toro, Panama. Sequencing was conducted in a single lane of Illumina HiSeq400 (NCBI Accession #: SAMN12669945, SAMN13924705-SAMN13924707) via the *de novo* approach Trinity software (Haas et al., 2013). BLASTx against the Metazoa (TaxID: 33208) database (e-value cutoff: e^-05^) was performed on the Super-Transcripts to add gene descriptions to each of the sequences. Key terms (e.g., Sirtuin, telomerase and telomere, POU, etc.) were used to find genes of interest among the BLAST descriptions (i.e. protein assignments).

### Expression normalization and profiling

The RNA-seq libraries of *Turritopsis dohrnii* life cycle stages were trimmed using Phred score cutoff of 10 as recommended for gene profiling analyses (MacManes, 2014) and aligned to the super-transcriptome using RSEM (Li and Dewey, 2011) to generate gene-level count data. The maSigPro Bioconductor package (Nueda et al., 2014) was used to perform normalization of the expression data using the upper quartile method (75% quantile). The data was analyzed in the reverse development sequence: colonial polyp to medusa, to cyst, to reversed polyp. Normalized expression values for each super-transcript were averaged per life cycle stage. Only genes with expression data consistent across biological stage replicates (i.e., expression present in all replicates or absent in all replicates) were used in further analyses and expression profiling visualization.

## Results and Discussion

### Expression profiling of candidate genes

Gene expression profiling of life cycle stages of *Turritopsis dohrnii* can reveal genetic networks underlying long lifespans, tissue regeneration, and cellular plasticity in metazoans. Super-transcripts have been reported to increase the accuracy of gene expression profiling analyses and provide distinct advantages over using a single representative sequence for a single gene, often the longest transcript isoforms (Davidson et al. 2017). In this work, we use super-transcripts constructed from *T. dohrnii’s* life cycle stages involved in reverse development (colonial polyp, medusa, cyst, reversed polyp) to profile the expression of Sirtuin (SIRT), telomere and telomerase-related genes, Heat-shock proteins (HSPs), and the Yamanaka Factor gene family (Oct, Sox, Klf, Myc). Only the expression profiles of super-transcripts with consistent expression among all biological replicates of *T. dohrnii* (i.e., present or absent in all replicates) are presented below.

#### Sirtuin proteins

Seven SIRT proteins have been identified in humans (SIRT1-SIRT7). Among those, the over-expression of Sir2 (silent information regulator 2, the invertebrate ortholog of SIRT2) has been linked to lifespan expansion in a variety of taxa such as yeast (Kaeberlein and Powers III 2007), nematodes (Tissenbaum and Guarente 2001), and fruit flies (Rogina and Helfand 2004). All seven SIRT proteins have been reported in chicken, mice, frogs, segmented worms, and sea anemones (specifically *Nematostella*), but only SIRT1, SIRT4, and SIRT6 have been identified in *Hydra* (Greiss and Gartner 2009).

Among the *T. dohrnii* super-transcripts, we identified 34 SIRT sequences with expectation value (e-value) e-^11^ to e-^128^ against the NR Metazoa database (Appendix A). We identified putative homologs for all seven SIRT proteins, SIRT6 being the most prevalent (Figure 1A). Notably, a wide range of metazoan taxa was identified as the top-hit species for the SIRT annotated sequences (Figure 1B-C). Some of these Sirtuins are present in *T. dohrnii* and *Nematostella*, but absent in *Hydra,* possibly due to the gene losses, a common occurrence in other metazoans (Greiss and Gartner 2009).

**Figure 1:**
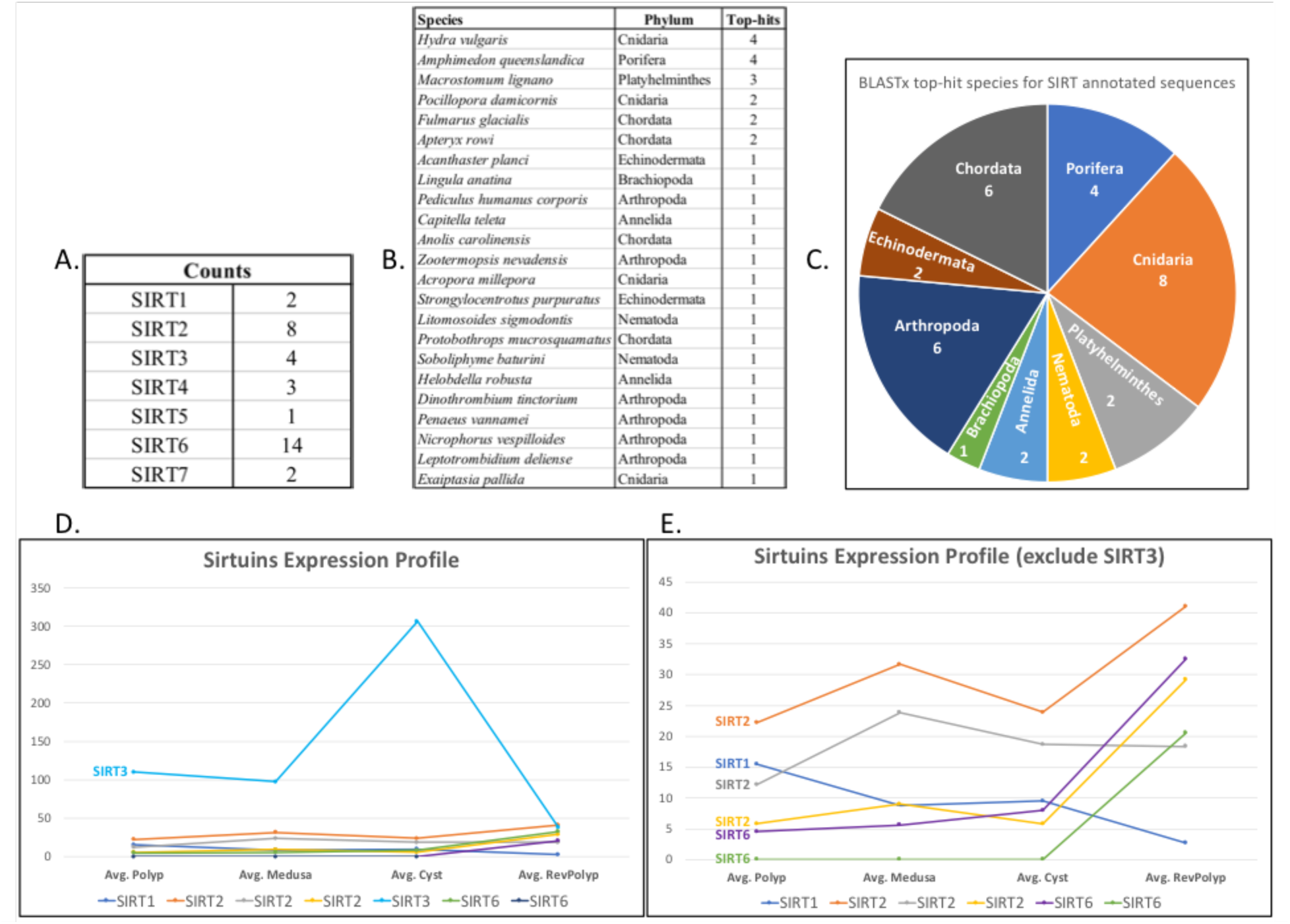
Sirtuin protein putative homologs in *T. dohrnii.* A) Number of super-transcripts for each SIRT sequence; B) Number of BLASTx top-hits for each species and their corresponding phylum; C) Pie chart of top-hit species for SIRT annotated sequences; D) Expression profiles of SIRT proteins (most consistent expression patterns across replicates chosen; complete list in Appendix A); E) Expression profiles of SIRT proteins excluding highest expressed SIRT3 sequence (D-light blue).

Among the 7 Sirtuins, SIRT1, 2, 3, and 6 show consistent expression among replicates and during the reverse development of *T. dohrnii* (Figure 1). SIRT3 is involved in mitochondrial energy metabolism, protects cells against oxidative stress, and has been reported to affect cellular lifespan and telomeres length (Lundberg et al. 2000), and is the only SIRT with experimental evidence directly correlated to human longevity (Bellizzi et al. 2005; Kincaid and Bossy-Wetzel 2013). It was also the highest expressed among all stages of *T. dohrnii* with a sharp peak in the cyst (Figure 1D, light blue). SIRT1, 2, and 6 expressions were low compared to SIRT 3 (Figure 1E, purple/green). SIRT6 (Figure 1E, yellow/orange) and SIRT 2 (Figure 1E, yellow/orange) sequences peaked in the reversed polyp. SIRT6 has significant implications in increasing the efficiency of DNA repair in long-lived mammals compared to their short-lived counterparts (Tian et al. 2019). In addition to DNA repair, SIRT6 protects telomeres and stabilizes the genome (Jia et al. 2012). SIRT2 is involved in cell-cycle control and development and is also known to be a marker of cellular senescence in humans (Grabowska et al. 2017). Lastly, SIRT1, which has a role in DNA repair, glucose metabolism, and insulin production and promotes cell survival in combination with delaying replicative senescence (Grabowska et al. 2017; Bellizzi et al. 2007), was the only SIRT in which transcriptional expression decreased from the colonial polyp to the reverse polyp stage (Figure 1E, blue).

### Telomeres and telomerase

The enzyme telomerase prevents the loss of genetic information during DNA replication via the elongation of protective sequences at the end of chromosomes. Except for epidermal cells, malignant and immortal germline cells, where indeterminate cell division occurs (Flores and Blasco 2010; Blasco 2007), telomerase is inactive in most somatic cells in metazoans. Previous work in *Turritopsis dohrnii* has shown that transcriptional regulation of genes involved in telomere maintenance occurs in the cyst stage (Matsumoto et al. 2019). In the presented work, we identified 56 telomere/telomerase-related super-transcripts (e-value ranges of e-^06^ – 0) (Appendix B). Many top hits corresponded to *Hydra*, other cnidarians, or sponges (Appendix B). Putative homologs of TEP1 (telomere-associated protein 1), a subunit of telomerase, were found to be the most numerous (Figure 2A).

**Figure 2:**
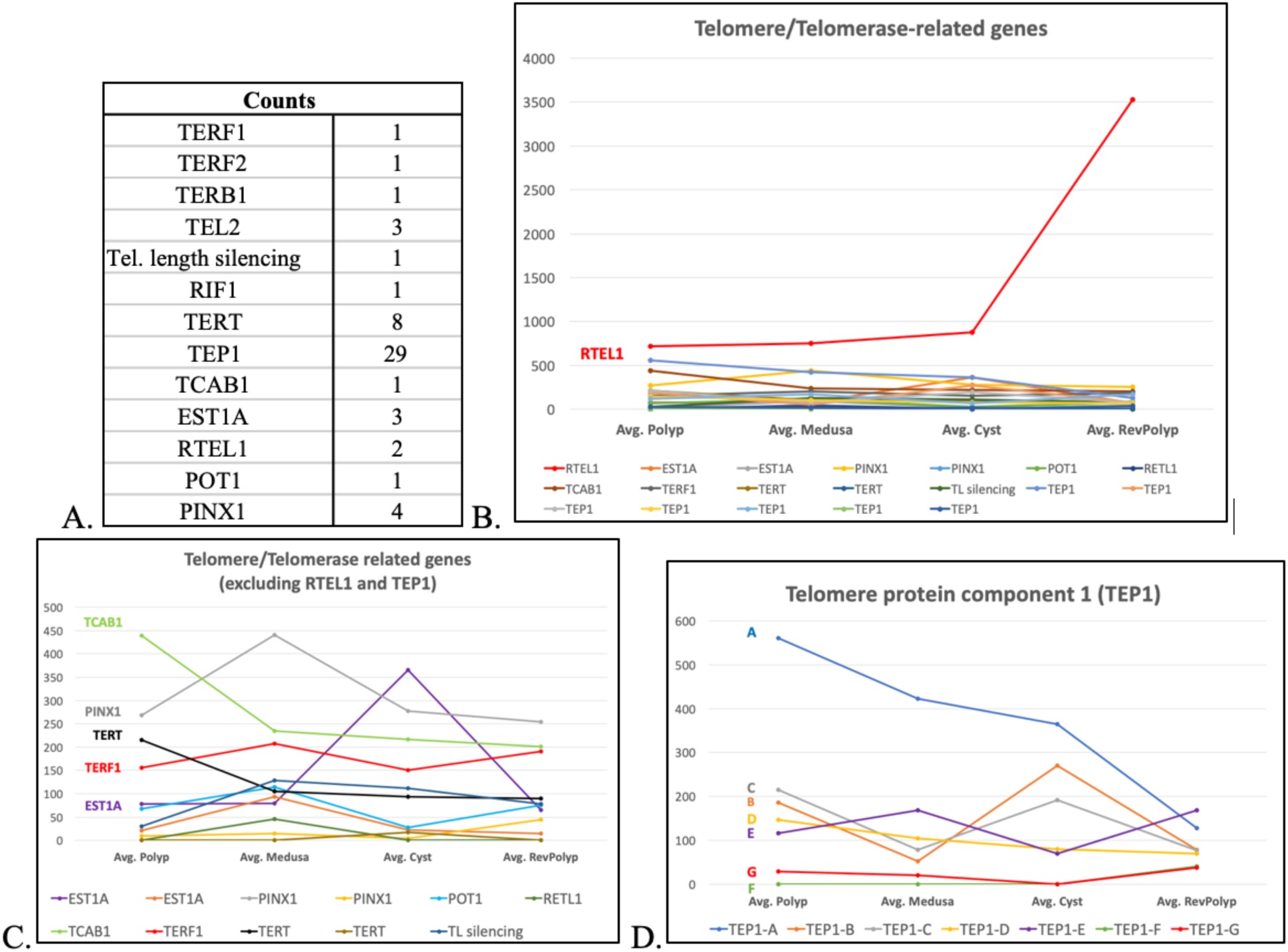
Telomere/Telomerase-related genes in *T. dohrnii*. A) Number of super-transcripts for each telomere/telomerase-related sequences; B) Expression profiles of all putative homologs (most consistent expression patterns across replicates chosen; complete list in Appendix B; C) Expression profiles excluding the RTEL1 and TEP1 sequences; D) Expression profiles of all TEP1 sequences.

Expression of the ’Regulator of telomere elongation 1’ (RTEL1), which plays a crucial role in regulating telomere length, DNA repair, and genomic stability in mice and humans (Barber et al. 2008; Uringa et al. 2011), was higher in *Turritopsis dohrnii*, with a sharp increase in the rejuvenated polyp (Figure 2B, red). An EST1A (telomerase-binding protein EST1A) sequence exhibited a sharp peak in expression at the cyst stage (Figure 2C, purple). EST1A binds and cooperates with TERT (telomerase reverse transcriptase) to elongate telomeres (Snow et al. 2003), and ectopic expression of the TERT alone has been shown to induce telomere lengthening in human cell cultures (Bodnar et al. 1998; Vaziri and Benchimol 1998). One TERT sequence showed only expression in the cyst (Figure 2C, brown). In contrast, the other TERT sequence had high expression in all stages, particularly in the colonial polyp (Figure 2C, black). Although more downstream work is necessary to determine the function of TERT and EST1A in *T. dohrnii*, our data indicate that the two genes are co-expressed in the cyst and may be working together to elongate telomeres before rejuvenation into the earlier polyp stage.

TEP1 (Telomerase Associated Protein 1), canonically active in immortalized murine cell lines but absent in somatic cells (Harrington et al. 1997), was the most abundant in this category, exhibiting a variety of expression profiles, with the highest expression in the colonial polyp stage (Figure 2D, blue). Similarly, TCAB1(Telomerase Cajal body protein 1), which is associated with the accumulation of cajal bodies that deliver RNA sequences to telomerase to elongate and maintain telomere length (Venteicher and Artandi 2009), was expressed among all stages but with highest expression in the colonial polyp (Figure 2C, light green).

Lastly, there was decreased expression of telomerase inhibitors TERF1 (Telomere repeat-binding factor 1) and PINX1 (PIN2/TERF-interacting telomerase inhibitor 1) during the transition from the medusa to the cyst (Figure 2C, light gray/ red). PINX1 has been reported to be a potent telomerase inhibitor (Zhou and Lu 2001), and TERF1 is known to bind and negatively regulate the length of telomeres and cooperates in the shelterin complex to protect chromosomal ends in mammals (Derevyanko et al. 2017). Combined with the previous reports of telomerase expression in the cyst (Matsumoto et al., 2019), our data indicate that telomerase may be involved in *T. dohrnii* ontogeny reversal.

### Heat-shock protein 70 and 90

Heat-shock proteins (HSPs) are activated in response to internal and external cellular stress and shock and promote quality control of protein folding processing, preventing misfolding and repairing or disposing of impaired proteins (i.e., autophagy) (Tower 2011). These molecular chaperones are ubiquitous, present in all domains of life, and have been extensively manipulated in invertebrate models to increase lifespan (Dubrez et al. 2019). As animals age, the production of HSPs and other stress-resistance factors slows down (Hou et al. 2010).

Experimental induction of HSPs can increase longevity in aged subjects by counteracting the accumulation of cellular damage that leads to physiological degeneration, malfunction, and disease (Murshid et al. 2013). Within the family, HSP90 and HSP70are the central facilitators of relieving cellular stress and shock (Calderwood 2007).

79 HSP70 and 49 HSP90 annotated super-transcripts were found in *T. dohrnii*, e-values ranging from e-^06^– 0 (Appendix C). Interestingly, *Hydra* was not the species with the most top hits and instead belonged to *Seriola lalandi dorsalis*, a ray-finned fish (Figure 3A). There were more chordate taxa, specifically fish, than cnidarians that correlated to top-hits in HSP putative homologs (Figure 3B). This may indicate that *T. dohrnii* utilizes some HSPs similar to vertebrate taxa that other Cnidaria do not possess and could be unique to the species.

**Figure 3:**
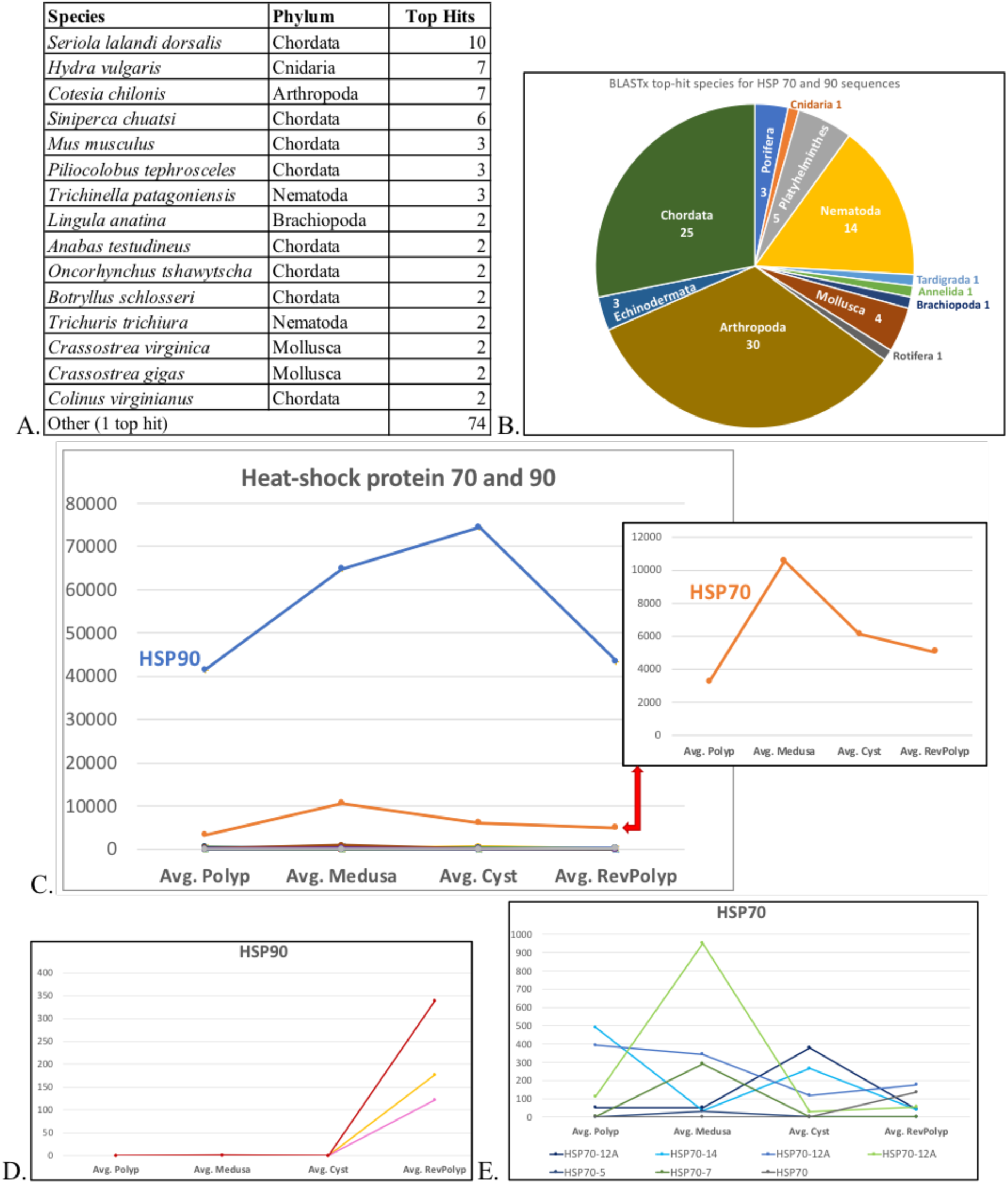
HSP70 and 90 in *T. dohrnii*. A) Number of BLASTx top-hits for each species and their corresponding phylum (all species with only one hit grouped as ’Other’); B) Pie chart of top species for HSP70 and 90 annotated sequences; C) Expression profiles of highest expressed HSP70 and 90 (most consistent expression patterns across replicates chosen; complete list in Appendix C); D) Expression profiles of HSP90 found only active in the reversed polyp; E) Expression profiles of other HSP70 found among *T. dohrnii* stages.

Both HSP90 and HSP70 were active in all stages of *T. dohrnii* (Figure 3). The most active HSP90 top-blast hit belonged to *Hydra,* while HSP70 belonged to *Rattus norvegicus*, the brown rat (Figure 3C, Appendix C). The HSP90 peaked at the cyst before dropping in the reversed polyp (Figure 3C, blue). The expression of HSP70 peaked in the medusa and decreased in the cyst and the reversed polyp (Figure 3C, orange). In addition, three HSP90 super-transcripts were absent or minimally expressed among all stages except for the reversed polyp (Figure 3D). The expression of other HSP70 sequences was varied, with some peaking in the cyst (Figure 3D black, light blue) and some in the medusa stage (Figure 3D light green, dark blue). HSP90 acts as a suppressor of the ubiquitin-proteasome system (UPS), while another HPS70 sequence found to be most active in the medusa and cyst stage (Figure 3E), acts as an activator of the UPS (Pratt et al. 2010). The two HSPs (HPS90 and HPS70) thus cooperate to regulate the UPS, where damaged or misfolded proteins are tagged for proteasomal degradation through ubiquitination (Löw 2011). The UPS plays a significant role in slowing aging, combating stress and damage, and promoting cellular/tissue regeneration in various metazoans (Löw 2011). Additionally, ubiquitin-related genes were also found to be differentially expressed and enriched in the cyst in a previous transcriptomic study of reverse development in *T. dohrnii* (Matsumoto and Miglietta 2021).

### Yamanaka transcription factors

Oct4 (POU5), Sox2, Klf4, and c-Myc, known as the Yamanaka Transcription Factors, have been shown to induce pluripotency in mammals (Takahashi et al. 2007; Takahashi and Yamanaka 2006), with POU5 and Sox2 considered essential factors that work cooperatively to induce cellular pluripotency, and Klf4 and c-Myc considered interchangeable (Takahashi et al. 2007; Takahashi and Yamanaka 2006). The gene families that each of the Yamanaka Factors resides in, namely POU, Sox, Klf, and Myc, have various roles in maintaining cellular pluripotency and development in metazoans, often modulated during disease and implicated in biomedical and regenerative research (Lefebvre et al. 2007; Gold et al. 2014; Zhang et al. 2020; Dang 1999). POU5, which Oct4 resides in, has only been reported in vertebrates, whereas other POU classes have undergone diversification before the emergence of the Eumetazoa (Gold et al., 2014). Sox2 and c-Myc, on the other hand, are highly conserved among Metazoa and are found universally among vertebrate and invertebrate taxa (Kondoh and Lovell-Badge 2015; Sarid et al. 1987). Klf4 has yet to be reported in Cnidaria (Steele et al. 2011) but has been found among other invertebrates such as roundworms (Ma et al. 2014; Hsieh et al. 2017).

We first conducted BLAST and alignment-based analyses on the annotated transcriptome and RNA-seq libraries of *T. dohrnii* published in (Matsumoto and Miglietta 2021) (summary in Table 1, full details of analyses in Appendix D). We also included the Thompson Factors (Lin28, Nanog), which are replaceable factors of Klf4 and c-Myc (Yu et al. 2007) to induce pluripotency in mammalian cells. This is the first step to assess whether the genetic network that enables cellular transdifferentiation in *T. dohrnii* involves pathways used in mammalian stem cells. Evidence from the *Hydra* genome has shown that c-Myc and Sox2 are the only factors present in this widely used model system (Chapman et al. 2010), bringing the authors to conclude that the stem cell genetic network in *Hydra* probably has an evolutionary origin independent from that used in mammalian stem cells. Understanding which of these factors are present in *T. dohrnii* is important to clarify the relationship between cnidarian stem cells and those of other animals, understand the evolution of pluripotency induction in Metazoa, and indicate to which extent *T. dohrnii* can be used as a system to study cellular plasticity and pluripotency at large.

**Table 1:**
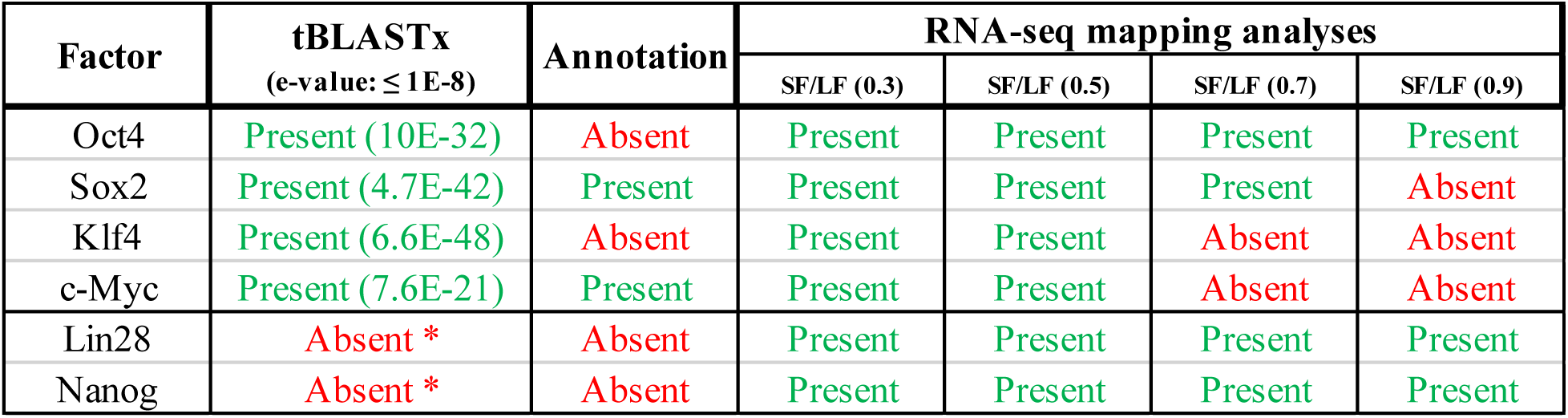
Summary of Yamanaka and Thomson transcription factors homology-based screening on published annotated transcriptome and RNA-seq libraries (Matsumoto and Miglietta, 2019). [*BLAST hit present under <400bps]; SF= Similarity fraction (minimum fraction of sequence identity between the read and reference), LF= length fraction (minimum length fraction that must match the reference sequence). Full details of analyses are shown in Appendix D.

#### Inference of homology using tBLASTx

We used all assembled transcripts filtered for biological contaminants (Matsumoto and Miglietta, 2021). All 12 query sequences, which included all variants of the six transcription factors reported in GenBank, had hits with regions of high-scoring pairs (Table 1; Appendix D). Only transcripts larger than 400 bp were annotated in Matsumoto and Miglietta (2021). Annotation results for tBLASTx hits against Oct4, Sox2, c-Myc, and Klf4 are reported below, while Lin28 and Nanog hits were shorter than 400bp and thus had no annotation (Appendix D; Matsumoto and Miglietta, 2021).

The presumed *Oct4* homolog in *Turritopsis dohrnii* (Td_DN89582_c0_g1_i1) identified in the reciprocal tBLASTx analyses above was annotated (i.e., protein assignment by BLASTx) as ’Brain-specific homeobox/POU domain protein 3’ with an e-value of 1.34e^-177^. The specific POU domain class 5 transcription factor was not found in the protein annotations (Appendix D). The presumed Sox2 homolog in *T. dohrnii* (Td_DN102764_c1_g1_i4) was annotated as the same protein, ’Sex determining region Y-box 2 protein’ with a e-value of 6.39e^-128^ (Appendix D). The presumed c-Myc (Myc proto-oncogene protein) homolog in *T. dohrnii* (Td_DN99862_c1_g1_i1) was also annotated as the same protein ’Myc proto-oncogene protein’, with a e-value of 2.55e^-23^ (Appendix D). The presumed *Klf4* homolog in *T. dohrnii* (Td_DN110215_c0_g1_i7) was annotated as ’Krueppel-like factor 5’ with a e-value of 3.48^e-47^. The specific *Klf4* factor was not found in among the annotations (Appendix D).

#### RNA-seq alignment analysis

We conducted RNA-seq mapping analyses using a range of length and similarity fractions (i.e., level of stringency) from 0.3 (least stringent) to 0.9 (most stringent). Read mapping of queried factors with lower stringency was performed due to the evolutionary distance between hydrozoans and humans. With a low-level stringency of a length fraction and similarity fraction of 0.3, all factors were recovered as present, and the consensus length of all of reads mapped to the query was similar to the actual length of all factors, merely 1-4 bp difference, except c-Myc with a 769 bp disparity (Appendix D). All factors were present with the stringency parameter of 0.3 and 0.5, but c-Myc and Klf4 were not present when the stringency parameter was set to 0.7. When a length and similarity fraction was set to 0.9, reads that map only to Oct4, Lin 28 and Nanog were recovered (Table 1 and Appendix D).

#### Expression profiling

We identified thirty super-transcripts annotated as POU, Sox, Klf, and Myc among the super-transcripts in *T. dohrnii* with e-values e-^08^– e-^168^ (Figure 4A, Appendix E). Most top hits belonged to cnidarian taxa (Appendix E).

**Figure 4:**
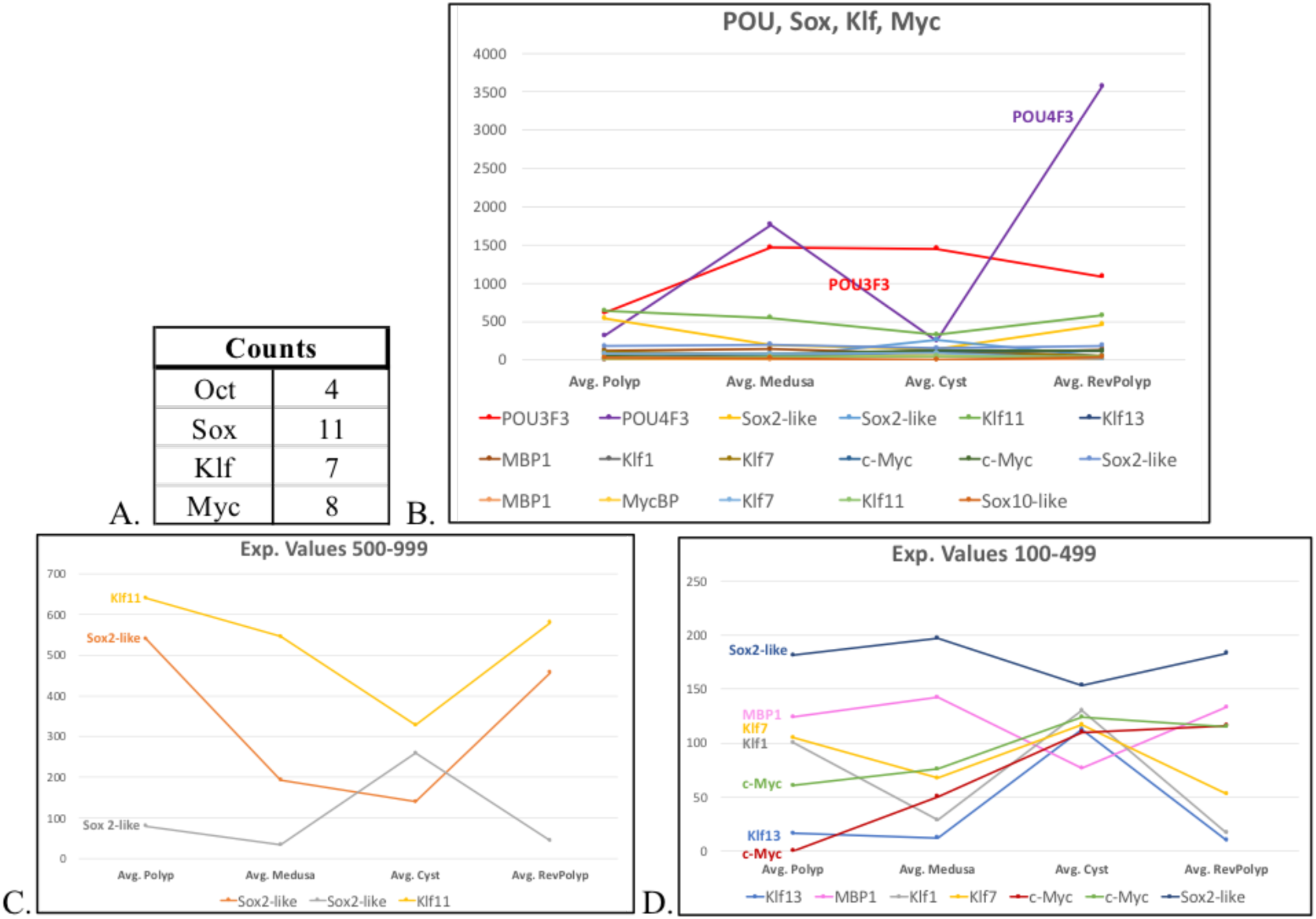
POU, Sox, Klf, and Myc in *T. dohrnii*. A) Number of super-transcripts for each gene family; B) Expression profiles of highest expressed POU, Sox, Klf, and Myc homologs (most consistent expression patterns across replicates chosen; complete list in Appendix E); C) Expression profiles of homologs in the normalized expression value range 500-999; D) Expression profiles of homologs in the normalized expression value range 100-499.

POU3F3 and POU4F3 homologs were active, with POU3F3’s expression peaking in the medusa and cyst, and POU4F3’expression enriched at the medusa and in the reversed polyp (Figure 4B, red/purple). POU3F3 is involved in neuronal development and works synergistically with Sox factors in mammals (Kuhlbrodt et al. 1998), while POU4F3 is a regulator of cell identity in the nervous system, and its dysregulation is central to hearing loss in mammals (Clough et al. 2004; Hertzano et al. 2004).

Genes in the Sox gene family are involved in cell fate determination during development (Lefebvre et al. 2007). Invertebrates have been reported to possess only a single Sox gene that is representative of the numerous Sox groups found in vertebrates (Bowles et al. 2000). The analyses, however, only consisted of *Drosophila*, roundworms, and an incomplete genome of the sea urchin, and thus may represent a partial view of the evolutionary history of Sox in invertebrates (Bowles et al. 2000). In *T. dohrnii,* we found Sox2 and Sox10-like putative homologs (Figure 4). Sox2-like homologs varied in expression profile (Figure 4CD), and the Sox2 factor was identified in the BLAST homology and alignment-based analyses (Table 2; Appendix D). The expression of Sox10-like was very low in all stages (Figure 4A, Appendix E).

We found homologs of Klf1, 7, 11, and 13 all active during *T. dohrnii’*s reversal (Figure 4). Klf11 was the most active, particularly in polyp stages (Figure 4C, yellow). The expression of Klf1, 7, and 13 increased from the medusa stage, peaked at the cyst, and declined after reversal (Figure 4D, yellow/gray). Klf factors are highly associated with the regulation of the cell cycle in mammals (Zhand et al., 2020; Tetreault et al., 2013). In nematodes, Klf factors (specifically Klf1, 2, and 3) are required for lifespan extension, and their deficiency reduces lifespan,and its over-expression increases longevity (Hsieh et al., 2017).

c-Myc is a proto-oncogene with diverse roles that include regulating the cell cycle, mediating cell lifespan, and maintaining pluripotent cell identity in mammals (Miller et al., 2012; Chappell et al., 2013). c-Myc was active during *T. dohrnii’s* reverse development along with its binding protein MBP (Figure 4D), which stimulates the activation of Myc (Taira et al. 1998). The expression of both c-Myc homologs found in *T. dohrnii* gradually increased through reverse development and peaked at the cyst and reversed polyp stages (Figure 4D, green/red).

## Conclusion

*T. dohrnii* offers a novel experimental paradigm to investigate *in vivo* the changes in gene activity that occur during reverse development and transdifferentiation. We profile the expressional changes during *T. dohrnii’s* reverse development of gene homologs of crucial tissue repair and regeneration regulators, inductors of the pluripotent state, and factors that promote longevity and increase lifespan. We show that *T. dohrnii* manipulates genetic networks that include SIRT3, POU factors, RTEL1, and HSP70/90, all of which are relevant genes in biomedical studies. *T. dohrnii* may represent a simple system to investigate regeneration, pluripotency, and longevity in Metazoa. Furthermore, our analyses indicate that the Yamanaka factors’ homologs may be present in *T. dohrnii.* Albeit not exhaustive, our analyses show a complex scenario, challenge our understanding of how the networks controlling a pluripotent cell state in Cnidaria relate to those of mammals, and identify areas of further exploration. They also highlight the need to expand the research on the genetics of induction of pluripotency to a variety of species that go beyond the classically used model systems.

