## Appendix for "The genetic networks of regeneration, cell plasticity, and longevity of the Immortal Jellyfish *Turritopsis dohrnii* (Cnidaria, Hydrozoa)": AppendixD_PreliminaryYFanalysis.docx

**Results**

Reported below are preliminary analyses on the published annotated transcriptome of *T. dohrnii* (Matsumoto and Miglietta, 2021) performed using BLAST and alignment-based approaches to provide insight on the potential presence of Yamanaka and Thomson factor homologs in *T. dohrnii*. Results of the Yamanaka and Thomson factors screening are summarized in Table 2. Understanding which of these factors are present in *T. dohrnii* is important as it may help us clarify the relationship between cnidarian stem cells and those of other animals, evolution of pluripotency induction in Metazoa, and indicate to which extent *T. dohrnii* can be used as a live model system to study cellular plasticity and pluripotency at large.

**Table**: **Summary of Yamanaka and Thomson transcription factors homology-based screening. [*BLAST hit present under <400bps] SF= Similarity fraction (minimum fraction of sequence identity between the read and reference), LF= length fraction (minimum length fraction that must match the reference sequence).**

| **Factor** | **tBLASTx**  **(e-value: ≤ 1E-8)** | **Annotation** | **RNA-seq mapping analyses** | | |  |
| --- | --- | --- | --- | --- | --- | --- |
|  |  |  | **SF/LF (0.3)** | **SF/LF (0.5)** | **SF/LF (0.7)** | **SF/LF (0.9)** |
| Oct4 | Present (10E-32) | Absent | Present | Present | Present | Present |
| Sox2 | Present (4.7E-42) | Present | Present | Present | Present | Absent |
| Klf4 | Present (6.6E-48) | Absent | Present | Present | Absent | Absent |
| c-Myc | Present (7.6E-21) | Present | Present | Present | Absent | Absent |
| Lin28 | Absent * | Absent | Present | Present | Present | Present |
| Nanog | Absent * | Absent | Present | Present | Present | Present |

*Inference of homology using BLAST*

For the reciprocal tBLASTx screening we used an e-value of ≤ 1e-^8^ and all mRNA variants of the *Homo sapiens* transcription factors available in GenBank were utilized. We also used all assembled transcripts that were filtered for biological contaminants, but not trimmed (i.e. 400bp or less removed), to avoid neglecting potential partial/incomplete transcripts (400bp or less) with significant alignments. All 12 query sequences, which included all variants of the six transcription factors reported in GenBank, had hits with regions of high scoring pairs (HSPs) (Table 2; Appendix J). Only transcripts larger than 400bp were kept in the final transcriptome assembly (Appendix A, Table 1A) and processed through the annotation pipeline (see details in Methods and Appendix L). Sequences larger than 400 bp were utilized for the subsequent annotation screening analyses. Lin28 and Nanog hits were shorter than 400bp and thus not further annotated (Appendix J). Annotation results for Oct4, Sox2, c-Myc, and Klf4 are reported below.

Transcript annotation screening analyses

The presumed Oct4 homolog in *T. dohrnii* (Td_DN89582_c0_g1_i1) identified in the reciprocal tBLASTx analyses above was annotated (i.e. protein assignment by BLASTx) as ‘Brain-specific homeobox/POU domain protein 3’ with a e-value of 1.34e^-177^. POU domain class 3, 6 and CF1A were represented, but the specific POU domain class 5 transcription factor was not found in the protein annotations (Appendix J). The presumed Sox2 homolog in *T. dohrnii* (Td_DN102764_c1_g1_i4) was annotated as the same protein, ‘Sex determining region Y-box 2 protein’ with a e-value of 6.39e^-128^ (Appendix J). The presumed c-Myc (Myc proto-oncogene protein) homolog in *T. dohrnii* (Td_DN99862_c1_g1_i1) was also annotated as the same protein ‘Myc proto-oncogene protein’, with a e-value of 2.55e^-23^ (Appendix J). The presumed Klf4 homolog in *T. dohrnii* (Td_DN110215_c0_g1_i7) was annotated as ‘Krueppel-like factor 5’ with a e-value of 3.48^e-47^ (Appendix J). Klf factors 1, 7, 8, 9, 10 and 11 were also found in the transcriptome, but Klf4 was not found in among the annotations (Appendix J).

*RNA-seq alignment analysis*

To further investigate the presence of the Yamanaka and Thomson factors, we conducted RNA-seq mapping analyses using a range of length and similarity fractions (i.e. level of stringency) from 0.3 (least stringent) to 0.9 (most stringent). Variants with the lowest e-value and highest bit-score larger than 400 bp was prioritized (i.e. Oct4 variant 1). Read mapping of queried factors with lower stringency was performed due to the evolutionary distance between hydrozoans and humans. With a low-level stringency of a length fraction and similarity fraction of 0.3, the presence of all factors were recovered, and the consensus length of all of reads mapped to the query was similar to the actual length of all factors, merely 1-4 bp difference, with the exception of c-Myc with a 769 bp disparity (Appendix J). All factors were present with the stringency parameter of 0.3 and 0.5, but c-Myc and Klf4 were not present when the stringency parameter was set to 0.7. When a length and similarity fraction was set to 0.9, reads that map only to Oct4, Lin 28 and Nanog were recovered.

**Discussion**

*Yamanaka Transcription Factor homologs in* T. dohrnii

A combination of four key transcription factors, Oct4, Sox2, Klf4 and c-Myc have been identified as the core transcription factors that govern the induction of pluripotency in murine (Takahashi and Yamanaka 2006) and human cells (Takahashi et al. 2007). Two additional transcription factors, Lin28 and Nanog, also known as the Thomson Factors, can replace Klf4 and c-Myc, and along with Oct4 and Sox2 can reprogram fully-differentiated cells into iPSC in humans (Yu and Silva 2008).

Only two of the genes, Sox2 and c-Myc, have been reported to be present in the phylum Cnidaria (Chapman et al. 2010; Jager et al. 2011; Millane et al. 2011). *Hydra* has been reported to possess 4 Myc homologues and the Sox B group (which includes the Sox2 gene) (Chapman et al. 2010). The fact that Oct4 and Klf4 were not detected in the *Hydra* genome prompted the conclusion that the stem cell genetic network in Hydrozoa has an independent evolutionary origin than of mammalian stem cells (Chapman et al., 2010).

Similar to *Hydra*, homologs of Oct4 and Klf4 were not detected in *T. dohrnii’*s transcriptome when looking at the annotation screening analysis only. However, the reciprocal BLAST and RNA-seq mapping analyses indicate that homologs of all Yamanaka factors, including Oct4 and Klf4, are recovered when SF/LF was set to 0.3 and 0.5 (Appendix J; Table 2). With more stringent SF/LF values, only Oct4 (SF/LF= 0.7) and Sox2 (SF/LF=0.7, 0.9) homologs were present. The Thomson factors, Nanog and Lin28, were recovered in the RNA-seq mapping analyses, regardless of SF/SL value, but were not found in the reciprocal BLAST and among protein annotations.

In summary, our analyses indicate that the Yamanaka factors' homologs may be present in *T. dohrnii*. Our results are different from what found in *Hydra*, where only homologs of Sox2 and Myc have been reported. They thus challenge our understanding of how the networks controlling a pluripotent cell state in Cnidaria relate to those of mammals. Albeit not exhaustive, our analyses show a complex scenario and an area in need of further exploration. They also highlight the need to expand the research on the genetics of induction of pluripotency to a variety of species that go beyond classically used model systems.

**Conclusion**

Lastly, we show evidence that supports the presence of the Yamanaka factors homologs (Oct4, Sox2, Klf4, c-Myc) in *T. dohrnii* of the Yamanaka factors homologs. This warrants for further exploration and more advanced ortholog analyses, to better understand the molecular control of cnidarian pluripotency and, more generally, the evolution of stem cells and their genetic networks in Metazoa*.*

**Methods**

Summary: Transcripts of interest were identified in our transcriptome using a multi-approach method: 1) tBlastx search; 2) RNA-seq analysis; 3) Annotation description screening analysis. For the tBlastx search, all variants of the human Yamanaka Transcription Factors were utilized (GenBank Accession found in Appendix L) and a e-value cutoff of e^-10^ was utilized. For the RNA-seq analysis, different stringencies of parameters (0.3, 0.5, 0.7 and 0.9 length and similarity fraction) were used with a mismatch cost value of 2 (linear gap cost), insertion value of 3 and deletion value of 3. The following key words that correlated to genes of interest were screened for and identified among BLAST annotations (case non-sensitive): Oct4- POU, Octamer, Oct; Sox2- Sox, Sex determining region Y; Klf4- Klf, Krueppel-like; c-Myc- Myc.

Method details, software and parameters:

**tBLASTx** (CLC-GUI)

- Database: Final transcriptome assembly
- Query: GenBank Accession (all human Yamanaka/Thompson factors and variants):
  - **Oct4** - NM_002701.5, NM_203289.5, NM_001173531.2, NM_203289.5
  - **Sox2**- NM_003106.3
  - **Klf4**- NM_004235.5, NM_001314052.1
  - **C-Myc**- NM_002467.6, NM_001354870.1
  - **Lin28**- NM_024674.5
  - **Nanog**- NM_024865.3, NM_001297698.1
- e-value cutoff: e^-10^
  - Word size: 6
  - Low complexity filter: true
  - Matrix: BLOSUM62 (default)
  - Gap cost: Existence 11, Extension 1 (default)
  - Max number of his sequences: 100

**RNA-seq mapping: CLC Genomic Workbench v8.0.1 mapper**

Toolbox > NGS Core Tools > Map Reads to Reference

- Input: Raw reads from 11 RNA-seq libraries
- Reference dataset: (GenBank Accession)
  - **Oct4** - NM_002701.5
  - **Sox2**- NM_003106.3
  - **Klf4**- NM_001314052.1
  - **C-Myc**- NM_002467.6
  - **Lin28**- NM_024674.5
  - **Nanog**- NM_024865.3
- Read alignment (all default except Length/Similarity fraction)
  - Mismatch cost- 2 (linear gap cost)
    - Insertion cost: 3
    - Deletion cost: 3
- Length fraction
  - Analysis 1- 0.3
  - Analysis 2- 0.5
  - Analysis 3- 0.7
  - Analysis 4- 0.9
- Similarity fraction
  - Analysis 1- 0.3
  - Analysis 2- 0.5
  - Analysis 3- 0.7
  - Analysis 4- 0.9
- Non-specific match handling- map randomly

**Annotation description screening**

- Key word search in transcriptome annotations (BLASTx protein assignments)
  - **Oct4**- POU, Octamer, Oct
  - **Sox2**- SOX, ‘Sex determining region Y’
  - **Klf4**- Klf, Krueppel-like
  - **c-Myc**- Myc
